# Whole exome sequencing of ENU-induced thrombosis modifier mutations in the mouse

**DOI:** 10.1101/174086

**Authors:** Kärt Tomberg, Randal J. Westrick, Emilee N. Kotnik, David R Siemieniak, Guojing Zhu, Thomas L. Saunders, Audrey C. Cleuren, David Ginsburg

**Author notes:** Department of Molecular Genetics and Genomics, Washington University in St. Louis, Missouri, United States of America. Corresponding author (DG).

## Abstract

Although the Factor V Leiden (FVL) gene variant is the most prevalent genetic risk factor for venous thrombosis, only 10% of FVL carriers will experience such an event in their lifetime. To identify potential FVL modifier genes contributing to this incomplete penetrance, we took advantage of a perinatal synthetic lethal thrombosis phenotype in mice homozygous for FVL (*F5^L/L^*) and haploinsufficient for tissue factor pathway inhibitor (*Tfpi*^+/-^) to perform a sensitized dominant ENU mutagenesis screen. Linkage analysis conducted in the 3 largest pedigrees generated from the surviving *F5^L/L^ Tfpi*^+/-^ mice (‘rescues’) using ENU-induced coding variants as genetic markers was unsuccessful in identifying major suppressor loci. Whole exome sequencing was applied to DNA from 107 rescue mice to identify candidate genes enriched for ENU mutations. A total of 3,481 potentially deleterious candidate ENU variants were identified in 2,984 genes. After correcting for gene size and multiple testing, *Arl6ip5* was identified as the most enriched gene, though not reaching genome-wide significance. Evaluation of CRISPR/Cas9 induced loss of function in the top 6 genes failed to demonstrate a clear rescue phenotype. However, a maternally inherited (not ENU-induced) *de novo* mutation (*Plcb4*^R335Q^) exhibited significant co-segregation with the rescue phenotype (p=0.003) in the corresponding pedigree. Thrombosis suppression by heterozygous *Plcb4* loss of function was confirmed through analysis of an independent, CRISPR/Cas9-induced *Plcb4* mutation (p=0.01).

**Author summary:** Abnormal blood clotting in veins (venous thrombosis) or arteries (arterial thrombosis) are major health problems, with venous thrombosis affecting approximately 1 in every thousand individuals annually in the United States. Susceptibility to venous thrombosis is governed by both genes and environment, with approximately 60% of the risk attributed to genetic influences. Though several genetic risk factors are known, >50% of genetic risk remains unexplained. Approximately 5% of people carry the most common known risk factor, Factor V Leiden. However, only 10% of these individuals will develop a blood clot in their lifetime. Mice carrying two copies of the Factor V Leiden mutation together with a mutation in a second gene called tissue factor pathway inhibitor develop fatal thrombosis shortly after birth. To identify genes that prevent this fatal thrombosis, we studied a large panel of mice carrying inactivating gene changes randomly distributed throughout the genome. We identified several genes as potential candidates to alter blood clotting balance in mice and humans with predisposition to thrombosis, and confirmed this protective function for DNA changes in one of these genes (*Plcb4*).

## Introduction

Venous thromboembolism (VTE) affects 1:1000 individuals in the US each year and is highly heritable [1, 2]. A single nucleotide variant (SNV) in the *F5* gene, referred to as Factor V Leiden (FVL, p.R506G) is present in 5-10% of Europeans, conferring a 2-4 fold increased risk for VTE [3]. Although ∼25% of VTE patients carry the FVL variant [4], only ∼10% of individuals heterozygous for FVL develop thrombosis in their lifetime.

To identify genetic variants that could potentially function as modifiers for FVL-associated VTE risk, we recently reported a dominant ENU screen [5] in mice sensitized for thrombosis. Mice homozygous for the FVL mutation (*F5^L/L^*) and haploinsufficient for tissue factor pathway inhibitor *(Tfpi^+/-^)* die of perinatal thrombosis [6]. After ENU mutagenesis, 98 G1 *F5^L/L^ Tfpi^+/-^* progeny survived to weaning (“rescues”) and 16 progeny exhibited successful transmission of the ENU-induced suppressor mutation. However, subsequent efforts to genetically map the corresponding suppressor loci were confounded by complex strain-specific differences introduced by the required genetic outcross [5]. Similar genetic background effects have complicated previous mapping efforts [7] and have been noted to significantly alter other phenotypes [8, 9]. Additional challenges of this mapping approach include the requirement for large pedigrees and limited mapping resolution, with candidate intervals typically harboring tens to hundreds of genes and multiple closely linked mutations.

High throughput sequencing methods have enabled the direct identification of ENU-induced mutations. Thus, mutation identification in ENU screens is no longer dependent upon an outcross strategy for gene mapping [10, 11]. We now report whole exome sequencing (WES) of 107 rescue mice (including 50 mice from the previously reported ENU screen [5]). Assuming loss of gene function as the mechanism of rescue, these WES data were analyzed gene-by-gene to identify genes enriched with mutations (mutation burden analysis). The *Arl6ip5* gene emerged as the top candidate suppressor locus from this analysis. However, an independent CRISPR/Cas9-generated *Arl5ip5* mutant allele failed to demonstrate highly penetrant rescue of the *F5^L/L^ Tfpi^+/-^* lethal phenotype. Surprisingly, a maternally inherited (not ENU-induced) *de novo* mutation (*Plcb4^R335Q^*) exhibited significant co-segregation with the rescue phenotype (p=0.003) in an expanded pedigree.

## Results and discussion

### Smaller rescue pedigrees on pure C57BL/6J background

In the previously reported ENU screen [5], viable *F5^L/L^ Tfpi^+/-^* rescue mice were outcrossed to the 129S1/SvImJ strain to introduce the genetic diversity required for subsequent mapping experiments. However, complex strain modifier gene interactions confounded this analysis and resulted in a large number of “phenocopies” (defined as viable *F5^L/L^ Tfpi^+/-^*mice lacking the original rescue mutation). To eliminate confounding effects of these thrombosis strain modifiers, we generated an additional 2,834 G1 offspring exclusively maintained on the C57BL/6J background. Fifteen new rescue pedigrees were established from this screen (S1 Table). The frequency, survival, weight, and sex distributions of identified rescues were consistent with our previous report (S1 Fig). Though many of the pedigrees previously generated on the mixed 129S1/SvImJ-C57BL/6J background generated >45 rescue progeny per pedigree (8/16) [5], all pedigrees on the pure C57BL/6J background yielded <36 rescue mice (most generating ≤5 rescues) (S1 Table). Significantly smaller pedigrees in comparison to the previous screen (p=0.010, S2 Fig) are likely explained by a generally positive effect of the hybrid 129S1/SvImJ-C57BL/6J strain background either directly on rescue fertility (hybrid vigor) or indirectly by reducing the severity of the *F5^L/L^* phenotype. The C57BL/6J and 129S1/SvImJ strains have been shown to exhibit significant differences in a number of hemostasis-related parameters, including platelet count and TFPI and tissue factor expression levels [12], with the genetic variations underlying such strain specific differences likely contributing to the genetic mapping complexity noted in the previous report [5].

### Linkage analysis using coding ENU variants fails to map suppressor loci

As the rescue pedigrees were maintained on a pure C57BL/6J background, the only genetic markers that could be used for mapping were ENU-induced variants. WES of one G1 or G2 member of the three largest pedigrees (1, 6, and 13, S2 Table), identified a total of 86 candidate ENU variants that were also validated by Sanger sequencing analysis (S3 Table). Of these 86 candidate genes, 69 were present in the G1 rescue but not its parents (G0), indicating that they were likely ENU-induced variants. These 69 variants were then further genotyped in all other rescue progeny in the respective pedigrees. Given the low number of identified genetic markers (20-26 per pedigree), these three pedigrees were poorly powered (29.6%, 21.7% and 39.4%, respectively) to identify the rescue variants by linkage analysis (S3-S5 Figs A). None of the 19 ENU variants tested in pedigree 1 (S3 Fig B), showed linkage with a LOD-score >1.5 (S3 Fig C). Similarly, 26 and 24 variants analyzed in pedigrees 6 and 13, respectively (S4, S5 Figs B) also failed to demonstrate a LOD-score >1.5 (S4, S5 Figs C). Failure to map the causal loci in any of these pedigrees was likely due to insufficient marker coverage. However, in these analyses, we could not exclude the contribution from a non-ENU-induced variant [13] or an unexpectedly high phenocopy rate. While WES has been successfully applied to identify causal ENU variants within inbred lines [14] and in mixed background lines [15, 16], whole genome sequencing (WGS) provides much denser and more even coverage of the entire genome (∼3,000 ENU variants/genome expected) and outperforms WES for mapping [11]. However, a WGS approach requires sequencing multiple pedigree members [10], or pooled samples at high coverage [11], resulting in considerably higher expense with current methods.

### WES identifies 6,771 ENU-induced variants in 107 rescues

In order to identify exonic ENU mutations, a total of 107 G1 rescues (57 from the current ENU screen and an additional 50 rescues with available material from the previous screen [5]), were subjected to WES (S2 Table). From ∼1.5 million initially called variants, 6,735 SNVs and 36 insertions-deletions (INDELs) within exonic regions were identified as potential ENU-induced mutations, using an in-house filtering pipeline (see Materials and methods). The most common exonic variants were nonsynonymous SNVs (47%), followed by mutations in 3’ and 5’ untranslated regions (31%) and synonymous SNVs (15%). The remaining variants (7%) were classified as splice site altering, stoploss, stopgain, or INDELs (Fig 1A). T/A -> C/G (47%), and T/A -> A/T (24%) SNVs were overrepresented, while C/G -> G/C (0.8%) changes were greatly underrepresented (Fig 1B), consistent with previously reported ENU studies [17, 18]. Since ENU is administered to the G0 father of G1 rescues, only female progeny are expected to carry induced mutations on the X chromosome, while males inherit their single X chromosome from the unmutagenized mother. Among the called variants, all chromosomes harbored a similar number of mutations in both sexes, with the exception of the X chromosome where a >35 fold increase in SNVs per mouse was observed in females (Fig 1C). The average number of exonic ENU mutations for G1 rescues was ∼65 SNV per mouse (Fig 1D), consistent with expected ENU mutation rates [10, 18]. These data suggest that most called variants are likely to be of ENU origin.

**Fig 1.**
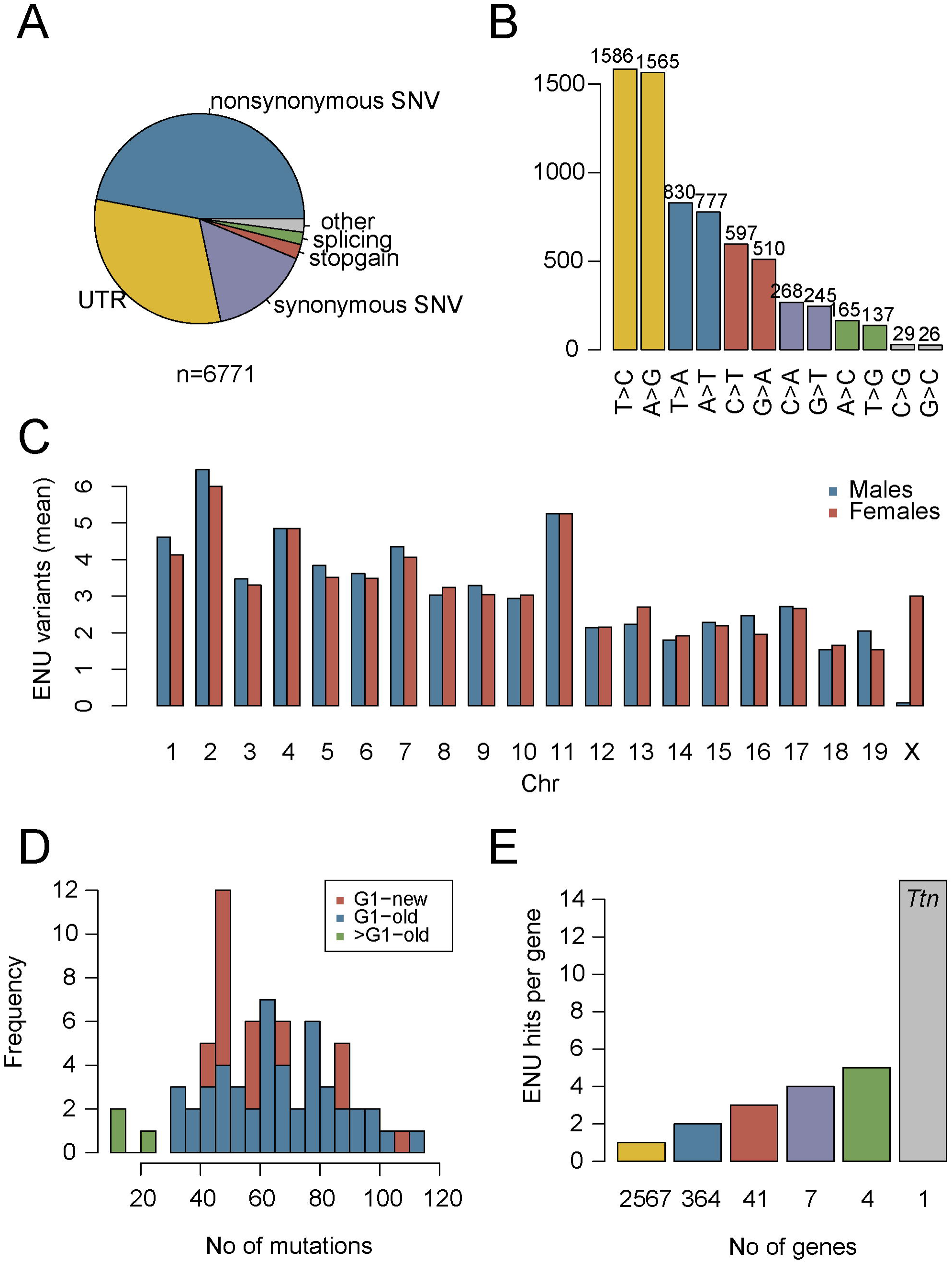
Distribution of ENU-induced mutations in WES data from 107 G1 rescues. A) Overview of mutation types for the 6,771 observed ENU-induced exonic variants. B) Distribution of missense mutations by nucleotide substitution type. C) Distribution of ENU-variants by chromosome. D) The average number of exonic SNVs is ∼65 for both the current (G1-new) and previous (G1-old) screen [5]. E) Number of genes (x-axis) sorted by the number of protein-altering ENU-induced mutations observed per gene (y-axis). Most genes (2,567) carry only 1 mutation. In contrast, the ∼0.1 megabase coding region of *Ttn* carries a total of 15 independent ENU variants.

### Mutation burden analysis identifies potential candidate thrombosis suppressor genes

WES data for 107 independent rescue mice were jointly analyzed to identify candidate genes that are enriched for potentially deleterious ENU-induced variants including missense, nonsense, frameshift, and splice site altering mutations (3,481 out of 6,771 variants in 2,984 genes, S4 Table). Similar mutation burden analyses have been used to identify genes underlying rare diseases caused by *de novo* loss-of-function variants in humans [19-22]. In our study, the majority of genes harbored only a single ENU-induced variant, with 15 SNVs identified in *Ttn*, the largest gene in the mouse genome (Fig 1E). After adjusting for coding region size and multiple testing (for 2,984 genes), the ENU-induced mutation burden of potentially deleterious variants was significantly greater than expected by chance for 3 genes (FDR<0.1, *Arl6ip5*, *Itgb6*, *C6*) and suggestive for 9 additional genes (FDR<0.25). Sanger sequencing validated 36 of the 37 variants in these 12 candidate genes (S4 Table). While in this study, stringent correction for multiple testing suggested no significant enrichment (*Arl6ip5* FDR=0.68, Fig 2), the potential power of this burden analysis is highly dependent on the number of possible genes that could result in a viable rescue. If there were 30 such genes in the genome and every one of the 107 rescue mice carried a mutation in one of these 30 genes, each gene would be, on average, represented by ∼3.5 mutations (107/30), with >7 genes expected to carry 5 or more mutations, which should have been sufficient to distinguish from the background mutation rate. However, if 500 genes could rescue the phenotype, sequencing close to a thousand mice would be required to achieve sufficient mapping power. The power could be further compromised by modifier genes with incomplete penetrance, imperfect predictions for potentially harmful mutations, and by the previously reported background survival rate for the rescue mice [6]. Due to the uncertainty of the power of these analyses, we proceeded to experimentally test the thrombosupressive effects of loss of function mutations in the genes identified by mutation burden analysis.

**Fig 2.**
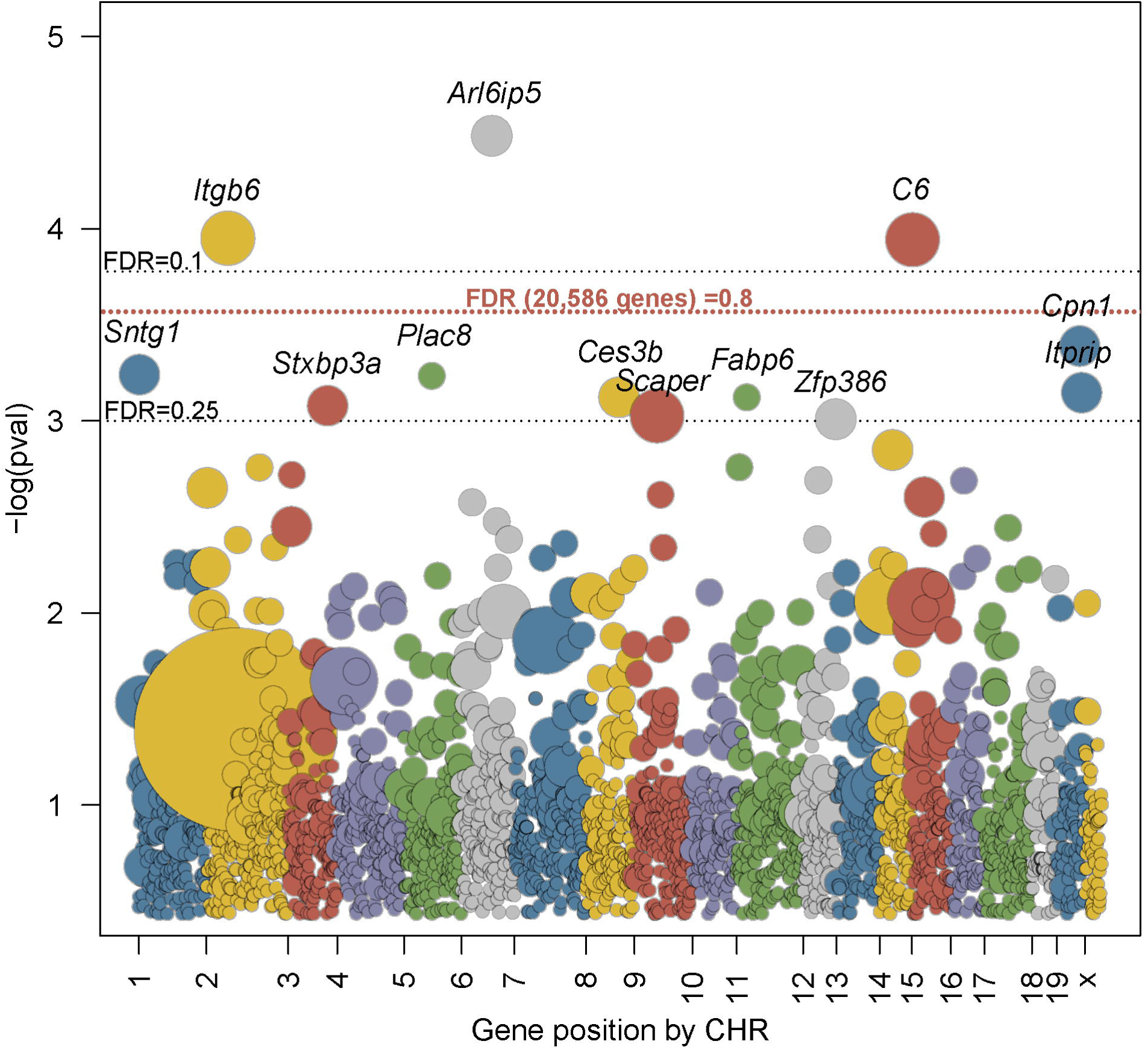
Mutation enrichment per gene in WES data from 107 G1 rescues. All genes with potentially deleterious ENU mutations are sorted by their chromosomal position on the x-axis, with the y-axis indicating the statistical significance (negative log of the p-value) of each gene’s enrichment based on 10^6^ permutations normalized to coding region size. Each dot represents a gene and the diameter is proportional to the number of mutations observed. Gray dotted lines represent FDR values of 0.1 and 0.25 (normalized to 2,984 genes carrying mutations). Red dotted line represents FDR value 0.8 from a more stringent test (normalized to all 20,586 genes in the simulation).

### Independent alleles for 6 candidate genes fail to replicate thrombosis suppression

Independent null alleles were generated with CRISPR/Cas9 for the top candidate genes (*Arl6ip5, C6, Itgb6, Cpn1, Sntg1 and Ces3b;* Fig 2) to test for thrombosuppression. From 294 microinjected zygotes with pooled guide RNAs targeting these 6 genes, we obtained 39 progeny. CRISPR/Cas9 genome editing was assessed by Sanger sequencing of the sgRNA target sites. Approximately 190 independent targeting events were observed across the 6 genes in 36 of the 39 mice including small INDELs, single nucleotide changes, and several large (>30bp) deletions or inversions. Targeted alleles were either homozygous, heterozygous, or mosaic, with the number of editing events varying greatly for different sgRNAs (2.5-85%). Two or more different CRISPR/Cas9-induced alleles for each of the candidate genes (S5 Table) were bred to isolation but maintained on the *F5^L^* background for subsequent test crossing. The progeny of *F5^L/L^ Tfpi^+/+^* mice crossed with *F5^L/+^ Tfpi^+/-^* mice (one of these parental mice also carrying the CRISPR/Cas9-induced allele) were monitored for survival of *F5^L/L^ Tfpi^+/-^* offspring (Table 1, S6 Table).

**Table 1.** Testing for rescue effect with CRISPR/Cas9-induced alleles.

| Gene | Total mice tested | No. of rescues w/o allele | No. of rescues with allele | P-value* |
| --- | --- | --- | --- | --- |
| *Arl6ip5* | 205 | 1 | 5 | 0.21 |
| *Itgb6* | 154 | 1 | 1 | 1 |
| *C6* | 106 | 0 | 0 | 1 |
| *Cpn1* | 139 | 0 | 1 | 1 |
| *Sntg1* | 223 | 3 | 4 | 1 |
| *Ces3b* | 219 | 2 | 1 | 1 |

\*Fisher's exact test

Over 100 progeny were generated for each of the candidate genes with no obvious rescue effect. A slight increase in rescues carrying *F5^L/L^ Tfpi^+/-^ Arl6ip5^+/-^* genotype was noted, although it remained non-significant after surveying 205 offspring (p=0.21, Table 1). Nonetheless, rescue of the *F5^L/L^ Tfpi^+/-^*phenotype by *Arl6ip5* haploinsufficiency cannot be excluded, particularly at reduced penetrance. Of note, rescue of *F5^L/L^ Tfpi^+/-^* lethality by haploinsufficiency for *F3* (the target of TFPI) only exhibits penetrance of ∼33% [5], a level of rescue which current observations cannot exclude for *Alr6ip5* and *Sntg1*. For most of the other candidate genes, the number of observed *F5^L/L^ Tfpi^+/-^* mice did not differ from the expected background survival rate for this genotype (∼2%) [6]. Though higher numbers of rescues were observed for offspring from the *Sntg1* cross, these were equally distributed between mice with and without the *Sntg1* loss-of-function allele.

### A *Plcb4* mutation co-segregates with the rescue phenotype in 3 G1 siblings and their rescue offspring

The number of G1 rescues produced from each ENU-treated G0 male is shown in Fig 3A. Though most of the 182 G0 males yielded few or no G1 rescue offspring, a single G0 produced 6 rescues out of a total of 39 offspring (Fig 3A), including the founder G1 rescue for the largest pedigree (number 13). This observation suggested a potential shared rescue variant rather than 6 independent rescue mutations from the same G0 founder. Similarly, another previously reported ENU screen identified 7 independent ENU pedigrees with an identical phenotype mapping to the same genetic locus, also hypothesized to result from a single shared mutation [7]. While rescue siblings could theoretically originate from the same mutagenized spermatogonial stem cell and share ∼50% of their induced mutations [23], such a common stem cell origin was excluded by exome sequence analysis in the rescue G1 sibs identified here (see Materials and methods).

**Fig 3.**
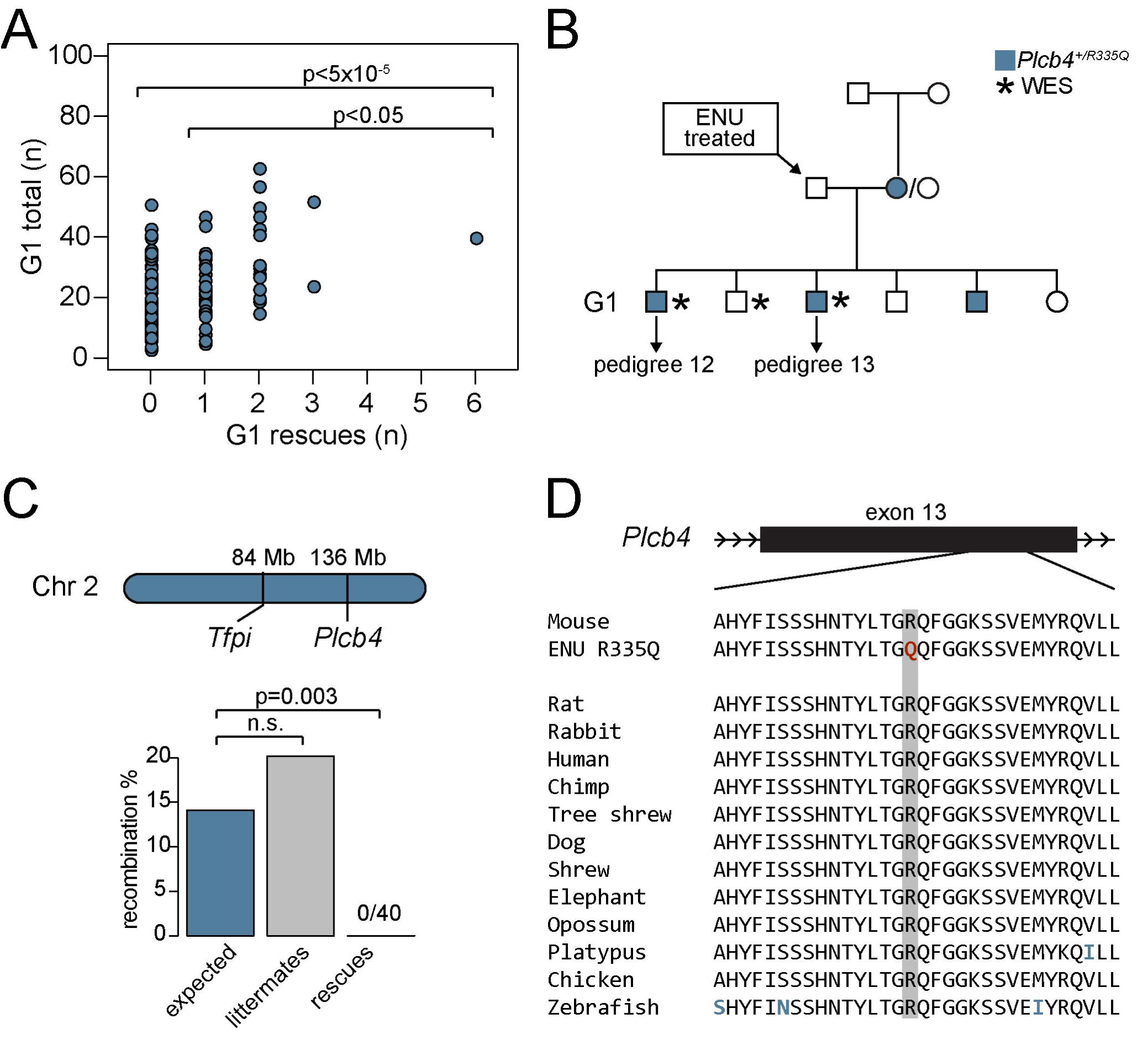
*Plcb4^R335Q^* co-segregates with the rescue phenotype in pedigrees 12 and 13. A) One ENU mating exhibited a significantly higher number of rescue progeny (n=6) compared to all ENU matings (p<5x10^-5^) and compared to ENU matings with ≥1 rescue progeny (p<0.05). B) One female in this ENU mating carried a *de novo* SNV (R335Q) in the *Plcb4* gene that was inherited in phase with the *Tfpi*^-^ allele that was inherited by 3 of the G1 rescues. C) The *Plcb4* gene is loosely linked to the *Tfpi* locus on chromosome 2, with a predicted recombination rate of 14.1%. No recombination was observed in 40 rescues from pedigree 12 and 13, while their littermates (n=149) exhibited close to the expected recombination rate. D) The *Plcb4^R335Q^* mutation lies in a highly conserved region in exon 13 (data from Multiz alignment on UCSC Genome Browser).

Analysis of WES for 3 of the G1 rescues originating from this common G0 founder male (Fig 3B, S2 Table) identified 3 protein-altering variants (*Plcb4^R335Q^*, *Pyhin1^G157T^*, and *Fignl2^G82S^*) shared among 2 or more of the 6 G1 rescues (S7 Table). *Plcb4^R335Q^* was detected as a *de novo* mutation in one of the non-mutagenized G0 females in phase with the *Tfpi* null allele (Fig 3B) and was present in 3 out of 6 G1 rescue siblings. *Plcb4* is located approximately 50 megabases upstream of the *Tfpi* locus on chromosome 2 (predicted recombination between *Plcb4* and *Tfpi* ∼14.1%) (Fig 3C) [24, 25]. While non-rescue littermates exhibited the expected rate of recombination between the *Plcb4^R335Q^* and *Tfpi* loci (20.2%), all 43 rescue mice (3 G1s and their 40 ≥G2 progeny) were non-recombinant and carried the *Plcb4^R335Q^* variant. This co-segregation between the *Plcb4^R335Q^* variant and the rescue phenotype is statistically significant (p=0.003; Fig 3C). *Plcb4^R335Q^*lies within a highly conserved region of *Plcb4* (Fig 3D) and is predicted to be deleterious by Polyphen-2 [26]. The other identified non-ENU variants (*Pyhin1^G157T^*and *Fignl2^G82S^*) did not segregate with the rescue phenotype (S6 Fig).

Although the estimated *de novo* mutation rate for inbred mice (∼5.4 x 10^-9^ bp/generation) is 200X lower than our ENU mutation rate, other *de novo* variants have coincidentally been identified in ENU screens [27]. Mutations identified by DNA sequencing of offspring from ENU screens will not distinguish between an ENU-induced and *de novo* origin, though the former is generally assumed, given its much higher prevalence in the setting of a mutagenesis screen. *De novo* mutations originating in the G0 paternal or maternal lineages will be identified by analysis of parental genotypes, as was the case for the *Plcb4^R355Q^* variant. However, this variant was originally removed from the candidate list by a filtering step based on the assumption that each ENU-induced mutation should be unique to a single G1 offspring. This filtering algorithm was very efficient for removing false positive variants in our screen and others [16]. However, our findings illustrate the risk for potential false negative results that this approach confers.

### Independent mutant allele for *Plcb4* recapitulates the rescue phenotype

An independent *Plcb4* null allele was generated by CRISPR/Cas9. Three distinct INDELs were identified by Sanger sequencing in the 25 progeny obtained from the CRISPR/Cas9-injected oocytes. One of these alleles introduced a single nucleotide insertion at amino acid 328, resulting in a frameshift in the protein coding sequence (*Plcb4^ins1^*, Fig 4A-B). A total of 169 progeny from a *F5^L/L^ Plcb4^+/ins1^*X *F5^L/+^ Tfpi^+/-^* cross yielded 11 *F5^L/L^ Tfpi^+/-^* rescue progeny surviving to weaning (Fig 4C, S8 Table). Ten of these 11 rescues carried the *Plcb4^ins1^* allele, consistent with significant rescue (p=0.01, Fig 4C) with reduced penetrance (∼40%). *Plcb4* encodes phospholipase C, beta 4 and has been recently associated with auriculocondylar syndrome in humans [28]. No role for PLCB4 in the regulation of hemostasis has been previously reported, and the underlying mechanism for suppression of the lethal *F5^L/L^ Tfpi^+/-^* phenotype is unknown.

**Fig 4.**
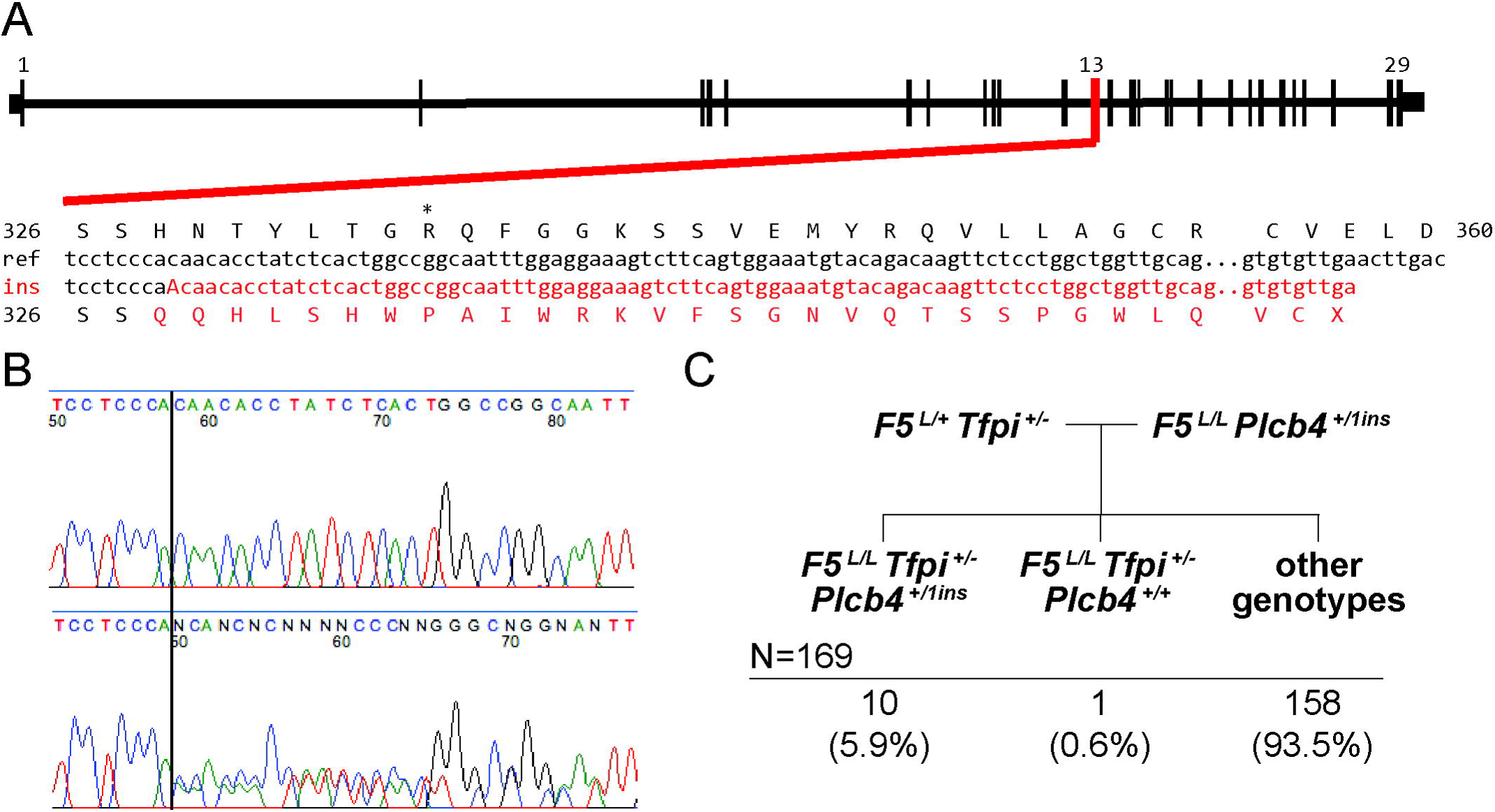
An independent CRISPR/Cas9 induced *Plcb4* allele validates the rescue phenotype. A) The CRISPR/Cas9-induced *Plcb4^ins1^*allele (insertion of the nucleotide ‘A’ at amino acid 328) results in a frameshift to the protein coding sequence leading to a premature stop codon. B) Sanger sequencing analysis of a wildtype mouse and a heterozygous mouse for the *Plcb4^ins1^* allele C) 169 progeny genotyped from a validation cross of *F5^L/L^ Plcb4^+/ins1^* mice with *F5^L/+^ Tfpi^+/-^*.

The above rescue of the *F5^L/L^ Tfpi^+/-^*phenotype by an independent *Plcb4* mutant allele, strongly supports the identification of the *de novo Plcb4^R355Q^* mutation as the causal suppressor variant for Pedigree 13. These findings are also most consistent with a loss-of-function mechanism of action for the *Plcb4^R355Q^*mutation. The lack of a positive signal from this genomic region by the linkage analysis described above (S5 Fig) is likely explained by the absence of a nearby genetically informative ENU variant (the closest, *Abca2* is located >50 Mb downstream from both *Tfpi* and *Plcb4* (S3 Table, S5 Fig)). Of note, 4 of the 107 rescue mice in the WES mutation burden analysis also carried a *Plcb4* mutation consistent with its suppressor function, though below the level of statistical significance. Nonetheless, these findings highlight the feasibility of our approach, given sufficient power.

In conclusion, we performed a dominant, sensitized ENU mutagenesis screen for modifiers of thrombosis. Analysis of extended pedigrees identified *Plcb4* as a novel thrombosis modifier. Though mutation burden analysis suggested several other potential modifier loci, including *Arl6ip5*, incomplete penetrance and the background phenocopy rate significantly limited the power to detect additional thrombosis suppressor genes. Future applications of this approach will likely require significantly larger sample sizes and/or a more stringent sensitized genotype for screening. Nonetheless, our findings demonstrate the power of a sensitized ENU screen and mutation burden analysis to identify novel loci contributing to the regulation of hemostatic balance and candidate modifier genes for thrombosis and bleeding risk in humans.

## Materials and methods

### Mice

Mice carrying the murine homolog of the FVL mutation [29] (*F5^L^*; also available from Jackson Laboratories stock #004080) or the TFPI Kunitz domain deletion (*Tfpi*^-^) [30] were genotyped using PCR assays with primers and conditions as previously described [29, 30], and maintained on the C57BL/6J background (Jackson Laboratories stock #000664). All animal care and procedures were performed in accordance with the Principles of Laboratory and Animal Care established by the National Society for Medical Research. The Institutional Animal Care and Use Committee at the University of Michigan has approved protocols PRO00005191 and PRO00007879 used for the current study and conforms to the standards of “The Guide for the Care and Use of Laboratory Animals” (Revised 2011).

### ENU screen

ENU mutagenesis was performed as previously described [5], with all mice on the C57BL/6J genetic background. Briefly, 189 *F5^L/L^* male mice (6-8 weeks old) were administered three weekly intraperitoneal injections of 90 mg/kg of ENU (N-ethyl-N-nitrosourea, Sigma-Aldrich). Eight weeks later, 182 surviving males were mated to *F5^L/+^ Tfpi^+/-^* females and their G1 progeny were genotyped at age 2-3 weeks to identify viable *F5^L/L^ Tfpi^+/-^* offspring (‘rescues’). *F5^L/L^ Tfpi^+/-^* G1 rescues were crossed to *F5^L/L^* mice on the C57BL/6J genetic background (backcrossed >20 generations) and transmission was considered positive with the presence of one or more rescue progeny. Theoretical mapping power in rescue pedigrees was estimated by 10,000 simulations using SIMLINK software [31].

### Whole exome sequencing

Gender, age, WES details, and other characteristics for 108 rescue mice are provided in S2 Table. Genomic DNA (gDNA) extracted from tail biopsies of 56 G1 offspring from the current ENU screen and from an additional 50 *F5*^L/L^ *Tfpi*^+/-^ mice on the C57BL/6J background from the previous screen [5] were subjected to WES at the Northwest Genomics Center, University of Washington. Sequencing libraries were prepared using the Roche NimbleGen exome capture system. DNA from an additional two rescue offspring was subjected to WES at Beijing Genomics Institute or Centrillion Genomics Technologies, respectively (S2 Table). These two libraries were prepared using the Agilent SureSelect capture system. 100 bp paired-end sequencing was performed for all 108 exome libraries using Illumina HiSeq 2000 or 4000 sequencing instruments. Two WES mice represented rescue pedigree 1: the G1 founder and a G2 rescue offspring. The latter was used for linkage analysis, but excluded from the burden analysis (S2 Table).

### WES data analysis

Average sequencing coverage, estimated by QualiMap software [32], was 77X, and >96% of the captured area was covered by at least 6 independent reads (S2 Table). All generated fastq files have been deposited to the NCBI Sequence Read Archive (Project accession number #PRJNA397141). A detailed description of variant calling as well as in-house developed scripts for variant filtration are online as a GitHub repository (github.com/tombergk/FVL_mod). In short, Burrows-Wheeler Aligner [33] was used to align reads to the *Mus Musculus* GRCm38 reference genome, Picard [34] to remove duplicates, and GATK [35] to call and filter the variants. Annovar software [36] was applied to annotate the variants using the Refseq database. All variants within our mouse cohort present in more than one rescue were declared non-ENU induced and therefore removed. Unique heterozygous variants with a minimum of 6X coverage were considered as potential ENU mutations. Among 107 whole exome sequenced G1 mice, 38 were siblings (13 sib-pairs and 4 trios, S2 Table). 190 heterozygous variants present in 2 or 3 mice (representing sibpairs or trios) out of 107 rescues were examined, with 15 found to be shared by siblings (S7 Table). Of the 7 sibs/trios sharing an otherwise novel variant, none shared >10% of their identified variants – inconsistent with the expected 50% for progeny originating from the same ENU-treated spermatogonial stem cell.

### Mutation frequency estimations

All ENU-induced variants predicted to be potentially harmful within protein coding sequences including missense, nonsense, splice site altering SNVs, and out-of-frame insertions-deletions (INDELs), were summed for every gene. The number of potentially damaging variants per gene was compared to a probability distribution of each gene being targeted by chance. Probability distributions were obtained by running 10 million random permutations using probabilities adjusted to the length of the protein coding region. A detailed pipeline for the permutation analysis is available online (github.com/tombergk/FVL_mod). Genes that harbored more potentially damaging ENU-induced variants than expected by chance were considered as candidate modifier genes. FDR statistical correction for multiple testing was applied as previously described [37].

### Variant validation by Sanger sequencing

All coding variants in pedigrees 1, 6, and 13 as well as all variants in candidate modifier genes from the burden analysis were assessed using Sanger sequencing. Variants were considered ENU-induced if identified in the G1 rescue but not its parents. All primers were designed using Primer3 software [38] and purchased from Integrated DNA Technologies. PCR was performed using GoTaq Green PCR Master Mix (Promega), visualized on 2% agarose gel, and purified using QIAquick Gel Extraction Kit (Qiagen). Sanger sequencing of purified PCR products was performed by the University of Michigan Sequencing Core. Outer primers were used to generate the PCR product which was then sequenced using the internal sequencing primers. All outer PCR primers (named: gene name+’_OF/OR’) and internal sequencing primers (named: gene name+’_IF/IR’) are listed in S9 Table.

### Guide RNA design and *in vitro* transcription

Guide RNA target sequences were designed with computational tools [39, 40] (http://www.broadinstitute.org/rnai/public/analysis-tools/sgrna-design or http://genome-engineering.org) and top predictions per each candidate gene were selected for functional testing (S10 Table). Single guide RNAs (sgRNA) for *C6*, *Ces3b*, *Itgb6*, and *Sntg1* were *in vitro* synthesized (MAXIscript T7 Kit, Thermo Fisher) from double stranded DNA templates by GeneArt gene synthesis service (Thermo Fisher) while the 4 sgRNAs for *Arl6ip5* were *in vitro* synthesized using the Guide-it sgRNA In Vitro Transcription Kit (Clontech). The sgRNAs were purified prior to activity testing (MEGAclear Transcription Clean-Up Kit, Thermo Fisher). Both the Wash and Elution Solutions of the MEGAclear Kit were pre-filtered with 0.02 μm size exclusion membrane filters (Anotop syringe filters, Whatman) to remove particulates from zygote microinjection solutions, thus preventing microinjection needle blockages.

### *in vitro* Cas9 DNA cleavage assay

Target DNA for the *in vitro* cleavage assays was PCR amplified from genomic DNA isolated from JM8.A3 C57BL/6N mouse embryonic stem (ES) cells [41] with candidate gene specific primers (S10 Table). *In vitro* digestion of target DNA was carried out by complexes of synthetic sgRNA and *S. pyogenes* Cas9 Nuclease (New England BioLabs) according to manufacturer’s recommendations. Agarose gel electrophoresis of the reaction products was used to identify sgRNA molecules that mediated template cleavage by Cas9 protein (S7 Fig). *Arl6ip5* was assayed separately, with one out-of-four tested sgRNAs successfully cleaving the PCR template.

### Cell culture DNA cleavage assay

Synthetic sgRNAs that targeted *Cpn1* were not identified by the *in vitro* Cas9 DNA cleavage assay. As an alternative assay, sgRNA target sequences (*Cpn1*-g1, *Cpn1*-g2) were cloned into plasmid pX330-U6-Chimeric_BB-CBh-hSpCas9 (Addgene.org Plasmid #42230) [42] and co-electroporated into JM8.A3 ES cells as previously described [43]. Briefly, 15 μg of a Cas9 plasmid and 5 μg of a PGK1-puro expression plasmid [44] were co-electroporated into 0.8x10^7^ ES cells. On days two and three after electroporation media containing 2 μg/ml puromycin was applied to the cells; then selection free media was applied for four days. Genomic DNA was purified from surviving ES cells. The *Cpn1* region targeted by the sgRNA was PCR amplified and tested for the presence of indel formation with a T7 endonuclease I assay according to the manufacturer’s instructions (New England Biolabs).

### Generation of CRISPR/Cas9 gene edited mice

CRISPR/Cas9 gene edited mice were generated in collaboration with the University of Michigan Transgenic Animal Model Core. A premixed solution containing 2.5 ng/μl of each sgRNA for *Arl6ip5*, *C6*, *Ces3b*, *Itgb6*, *Sntg1*, and 5 ng/μl of Cas9 mRNA (GeneArt CRISPR Nuclease mRNA, Thermo Fisher) was prepared in RNAse free microinjection buffer (10 mM Tris-Hcl, pH 7.4, 0.25 mM EDTA). The mixture also included 2.5 ng/μl of pX330-U6-Chimeric_BB-CBh-hSpCas9 plasmid containing guide *Cpn1*-g1 and a 2.5 ng/μl of pX330-U6-Chimeric_BB-CBh-hSpCas9 plasmid containing guide *Cpn1*-g2 targeting *Cpn1* (S10 Table). The mixture of sgRNAs, Cas9 mRNA, and plasmids was microinjected into the male pronucleus of fertilized mouse eggs obtained from the mating of stud males carrying the *F5*^L/+^ *Tfpi*^+/-^ genotype on the C57BL/6J background with superovulated C57BL/6J female mice. Microinjected eggs were transferred to pseudopregnant B6DF1 female mice (Jackson Laboratories stock #100006). DNA extracted from tail biopsies of offspring was genotyped for the presence of gene editing. The *Plcb4* allele was targeted in a separate experiment in collaboration with the University of Michigan Transgenic Animal Model Core using a pX330-U6-Chimeric_BB-CBh-hSpCas9 plasmid that contained guide *Plcb4* (5 ng/μl).

### CRISPR allele genotyping

Initially, sgRNA targeted loci were tested using PCR and Sanger sequencing (primer sequences provided in S10 Table). Small INDELs were deconvoluted from Sanger sequencing reads using TIDE software [45]. A selection of null alleles from >190 editing events were maintained for validation (S5 Table). Large (>30 bp) deletions were genotyped using PCR reactions that resulted in two visibly distinct PCR product sizes for the deletion and wildtype alleles. Expected product sizes and genotyping primers for each deletion are listed in S5 Table. All genotyping strategies were initially validated using Sanger sequencing.

### Picogreen DNA quantification and qPCR

A qPCR approach was applied to exclude large on-target CRISPR/Cas9-induced deletions. All DNA samples were quantified using the Quant-iT PicoGreen® dsDNA Assay Kit (Life Technologies) and analyzed on the Molecular Devices SpectraMax® M3 multi-mode microplate reader using SoftMax® Pro software and diluted to 5ng/μl. Primer pairs were designed for each gene using Primer Express 3.0 software (S9 Table) and samples were measured in triplicate using Power SYBR® Green PCR Master Mix (Thermo Fisher Scientific) on a 7900 HT Fast Real-Time PCR System (Applied Biosystems) with DNA from wildtype C57BL/6J mice used as a reference. While large CRISPR/Cas9 induced deletions extending the borders of the PCR primers have been reported [46], qPCR did not detect evidence for a large deletion in any of the CRISPR targeted genes.

### Statistical analysis

Kaplan-Meier survival curves and a log-rank test to estimate significant differences in mouse survival were performed using the ‘survival’ package in R [47]. A paired two-tailed Student’s t-test was applied to estimate differences in weights between rescue mice and their littermates. Fisher’s exact tests were applied to estimate deviations from expected proportions in mouse crosses. Mendelian segregation for CRISPR/Cas9-induced alleles among non-rescue littermates was assessed in a subset of mice by Sanger sequencing and then assumed for the rest of the littermates in the Fisher’s exact tests. Benjamini and Hochberg FDR for ENU burden analysis, Student’s t-tests, and Fisher’s exact tests were all performed using the ‘stats’ package in R software [48]. Linkage analysis was performed on the Mendel platform version 14.0 [49] and LOD scores ≥3.3 were considered genome-wide significant [50].

## Acknowledgments

We acknowledge Wanda Filipiak, Galina Gavrilina, Elizabeth Hughes, and Michael Zeidler for assistance in the preparation of CRISPR/Cas9 reagents and gene edited transgenic mice in the Transgenic Animal Model Core of the University of Michigan’s Biomedical Research Core Facilities. We thank the University of Michigan DNA Sequencing Core and the Northwest Genomics Center at the University of Washington, Department of Genome Sciences for sequencing services.

## Supporting information

**S1 Fig. A sensitized ENU suppressor screen for thrombosis modifiers**

A) The ENU screen strategy is depicted here, along with the total numbers of G1 offspring observed by genotype. B) Survival curves for G1 rescue mice. Approximately 50% of the rescue mice died by 6 weeks of age, with no significant survival difference observed between females and males (p=0.077), though females were underrepresented compared to males during the initial genotyping (28 females compared to 48 males, p=0.022). C-D) Weight at genotyping (at 14-21 days) was on average 25-30% smaller for G1 rescues than their littermates (p=7x10^-13^). E) Survival of rescue mice beyond G1 (≥G2) is also reduced, with worse outcome in females (p=0.002). Across all pedigrees, mice beyond G1 (≥G2) continued to exhibit reduced survival with more pronounced underrepresentation of females (p=0.002), and F) an average ∼22% lower body weight compared to littermates (mean defined as 100%) at the time of genotyping (p=2x10^-16^).

**S2 Fig. Size distribution of ENU pedigrees**

The ENU rescue pedigrees from the previous screen (n=16, [5]) are significantly larger than the ENU rescue pedigrees observed in the current screen (p=0.010, n=15, S1 Table).

**S3 Fig. Genetic mapping of ENU-induced variants in pedigree 1**

A) Overview of pedigree 1 (only rescue mice displayed). B) All coding ENU-induced mutations identified by WES were genotyped in all rescues from the pedigree by Sanger sequencing. Blue boxes indicate presence and red boxes indicate absence of the mutation. P1-P3 refers to 3 parental genotypes (G0 male and 2 untreated females). C) Linkage analysis using the ENU-induced variants from (B) as genetic markers.

**S4 Fig. Genetic mapping of ENU-induced variants in pedigree 6**

A) Overview of pedigree 6 (only rescue mice displayed). B) All coding ENU-induced mutations identified by WES were genotyped in most rescues from the pedigree by Sanger sequencing. Blue boxes indicate presence and red boxes indicate absence of the mutation. P1-P3 refers to 3 parental genotypes (G0 male and 2 untreated females). C) Linkage analysis using the ENU-induced variants from (B) as genetic markers.

**S5 Fig. Genetic mapping of ENU variants in pedigree 13**

A) Overview of pedigree 13 (only rescue mice displayed). B) All coding ENU-induced mutations identified by WES were genotyped in all rescues from the pedigree if present in key mice 3 and 5 by Sanger sequencing. Blue boxes indicate presence and red boxes indicate absence of the mutation. P1-P3 refers to 3 parental genotypes (G0 male and 2 untreated females). C) Linkage analysis using the ENU-induced variants from (B) as genetic markers.

**S6 Fig. Segregation analysis for *Pyhin1* and *Fignl2* in pedigree 13**

Segregation analysis in pedigree 13 for A) *Pyhin1* and B) *Fignl2* variants. Blue boxes indicate presence and red boxes indicate absence of the mutation. White boxes indicate untested mice, while light red boxes indicate untested mice with assumed absence of the mutation.

**S7 Fig. *In vitro* cleavage assay for sgRNAs**

A) sgRNA+Cas9 targeting created double strand breaks in DNA templates obtained from genomic DNA by PCR. Expected sizes after sgRNA+Cas9 endonuclease activity: 430bp/240bp (*Ces3b*), 334bp/273bp (*Sntg1*), 530bp/275bp (*Itgb6*), and 383bp/296bp (*C6*). B) sgRNA+Cas9 complexes targeting *Cpn1* using two different guides (g1, g2) failed to generate detectable double strand breaks. Positive control (P.C.) was added to ensure Cas9 protein activity, with expected sizes after cleavage (390bp/140bp) indicated by white stars.

**S1 Table. Overview of successfully progeny tested rescues**

**S2 Table. Rescue mice subjected to WES**

**S3 Table. Variants identified for pedigrees 1, 6, and 13**

**S4 Table. ENU-induced coding variants in WES data S5 Table. CRISPR/Cas9 induced alleles**

**S6 Table. Validation crosses with CRISPR/Cas9 induced alleles**

**S7 Table. Shared variants between 2-3 mice in WES data**

**S8 Table. *Plcb4^ins1^* validation cross**

**S9 Table. Primer sequences**

**S9 Table. Sequences and genotyping data for gRNAs**

