## Supplementary figures and images for "Whole exome sequencing of ENU-induced thrombosis modifier mutations in the mouse"

### Supplementary Materials

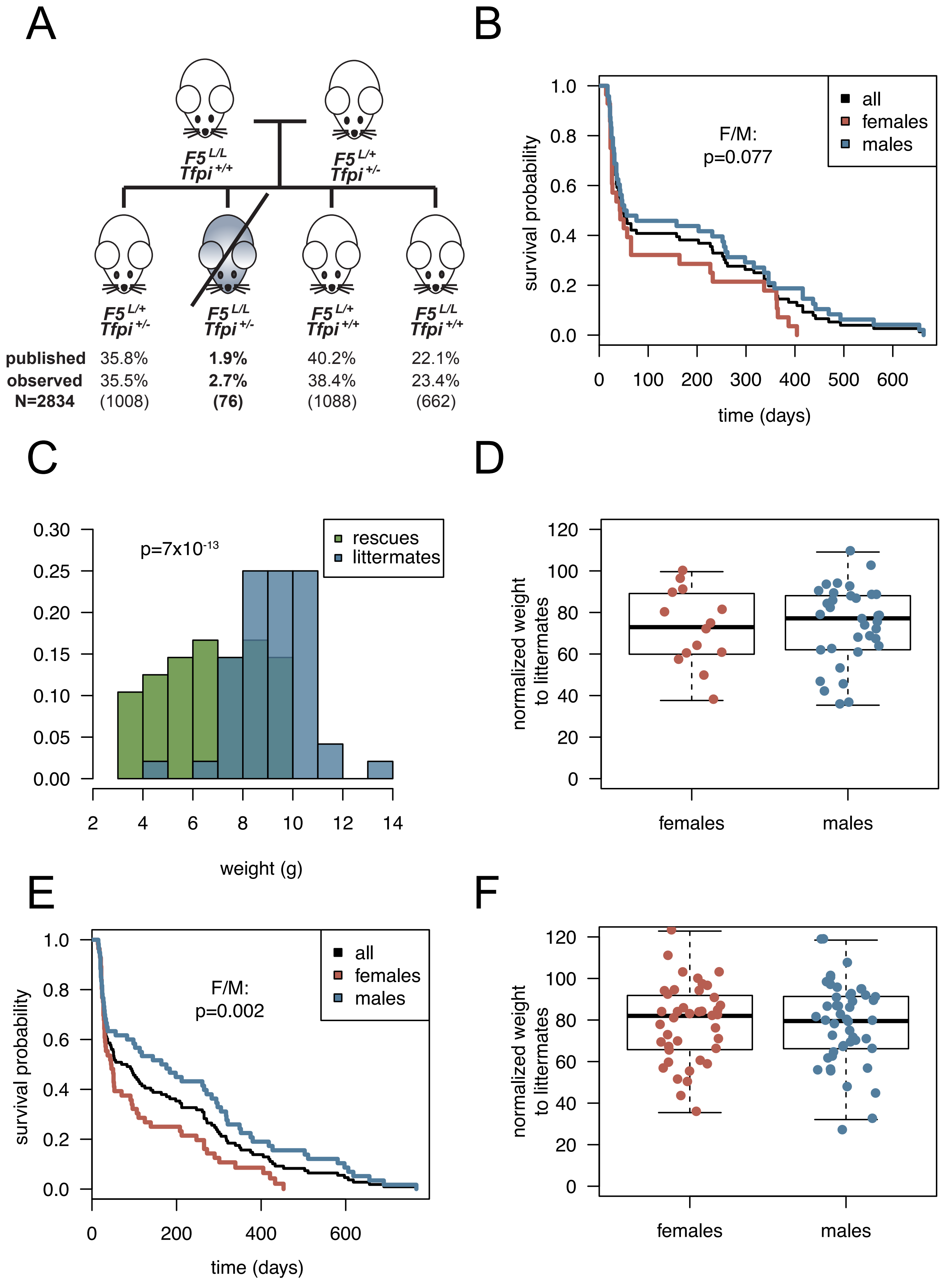

### Supplementary Materials

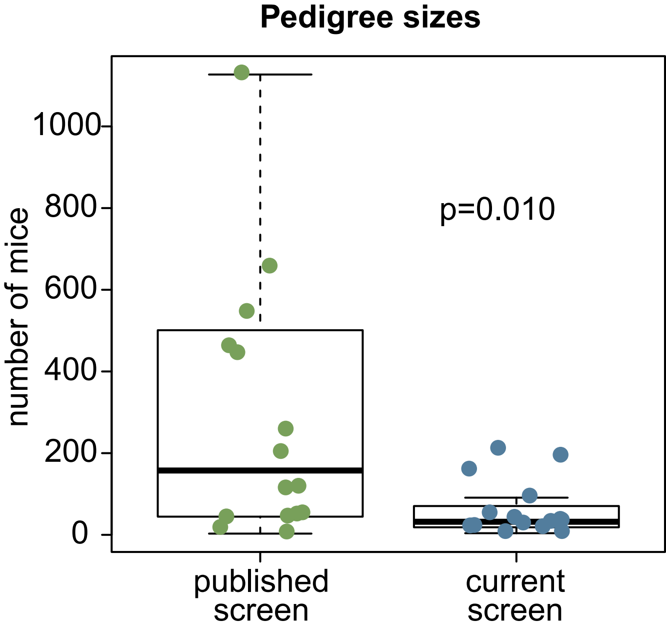

### Supplementary Materials

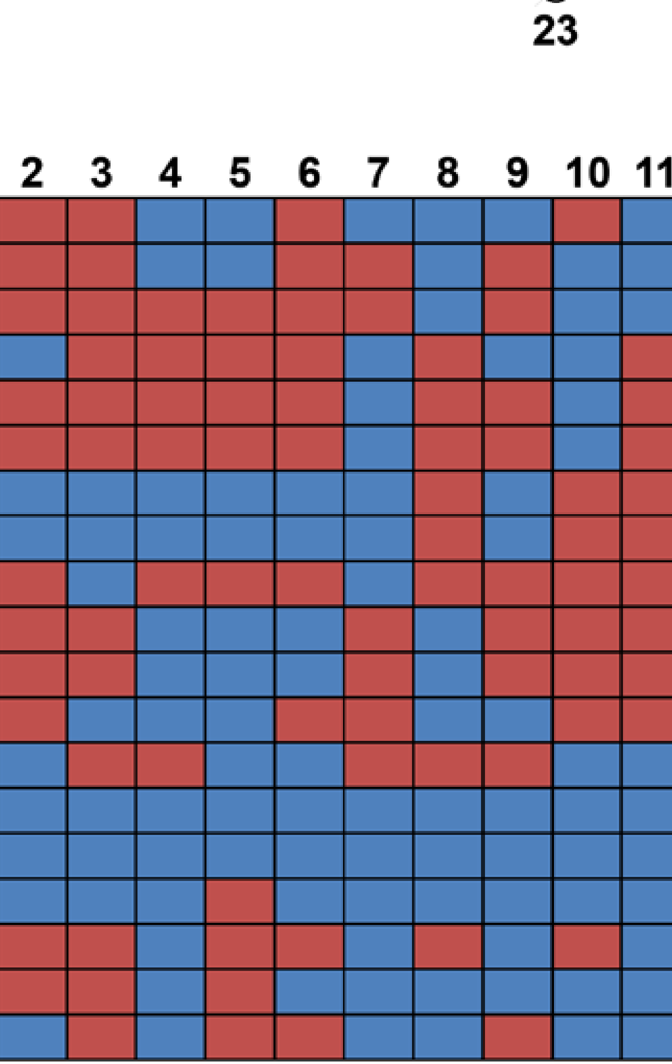

### Supplementary Materials

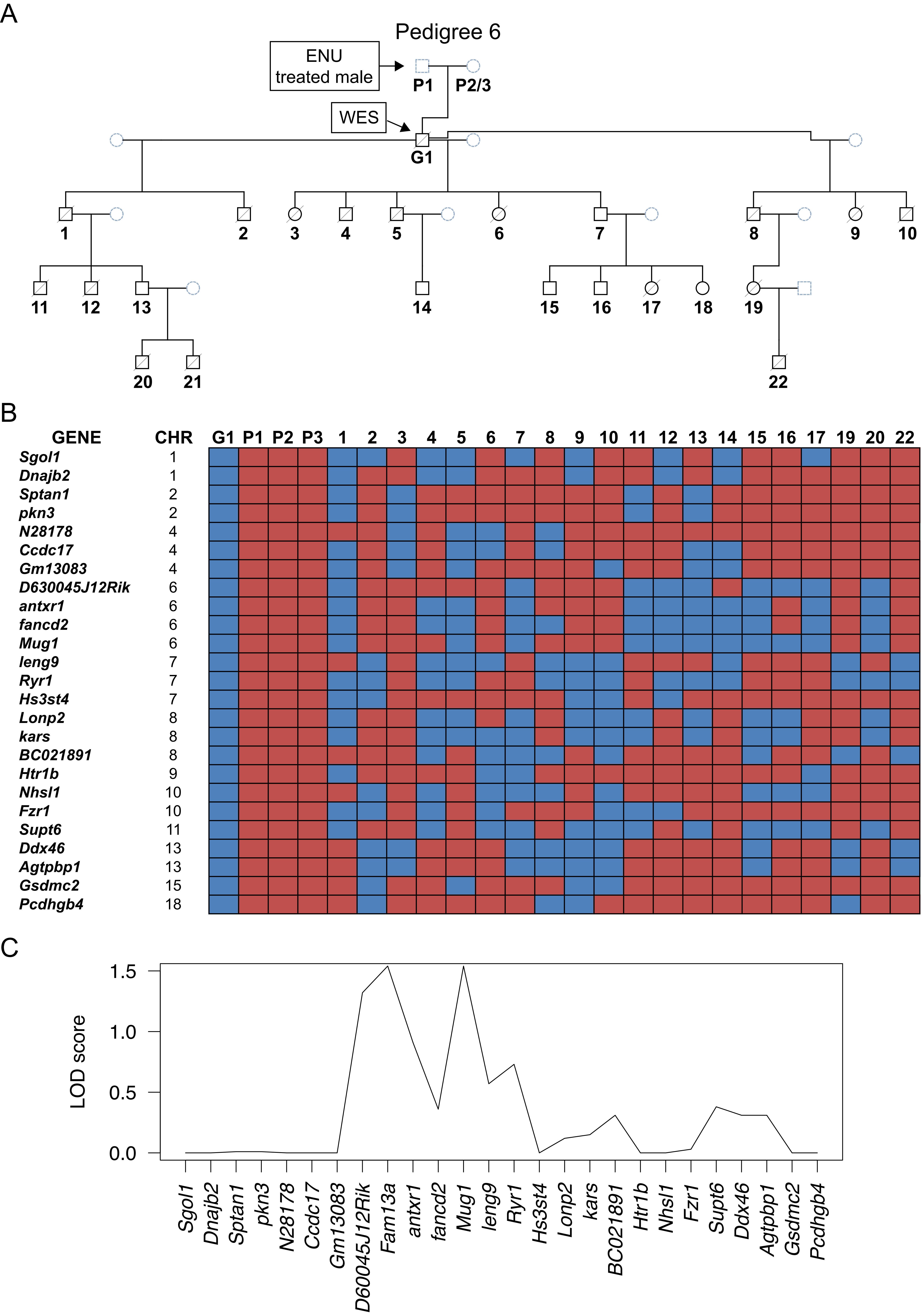

### Supplementary Materials

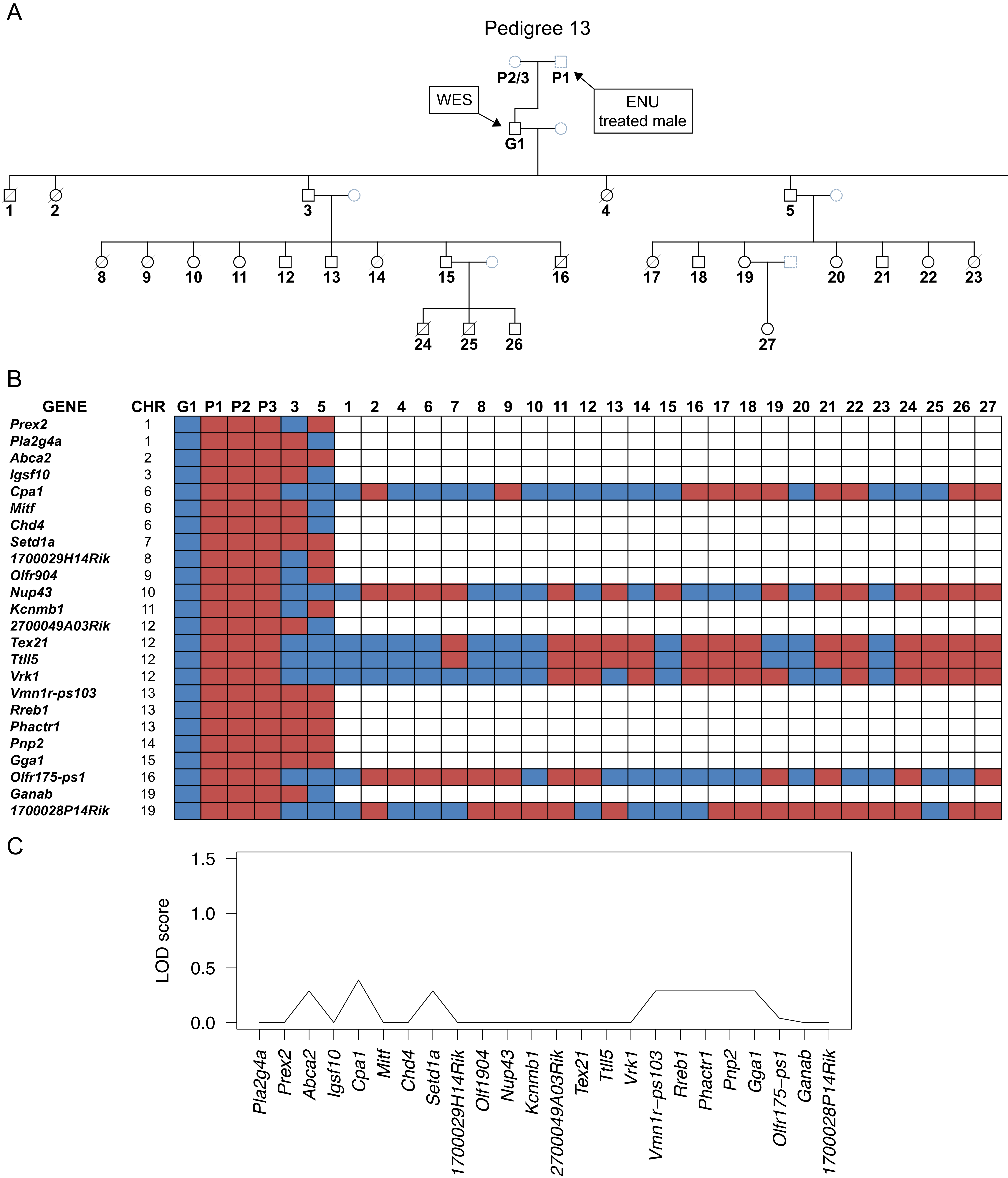

### Supplementary Materials

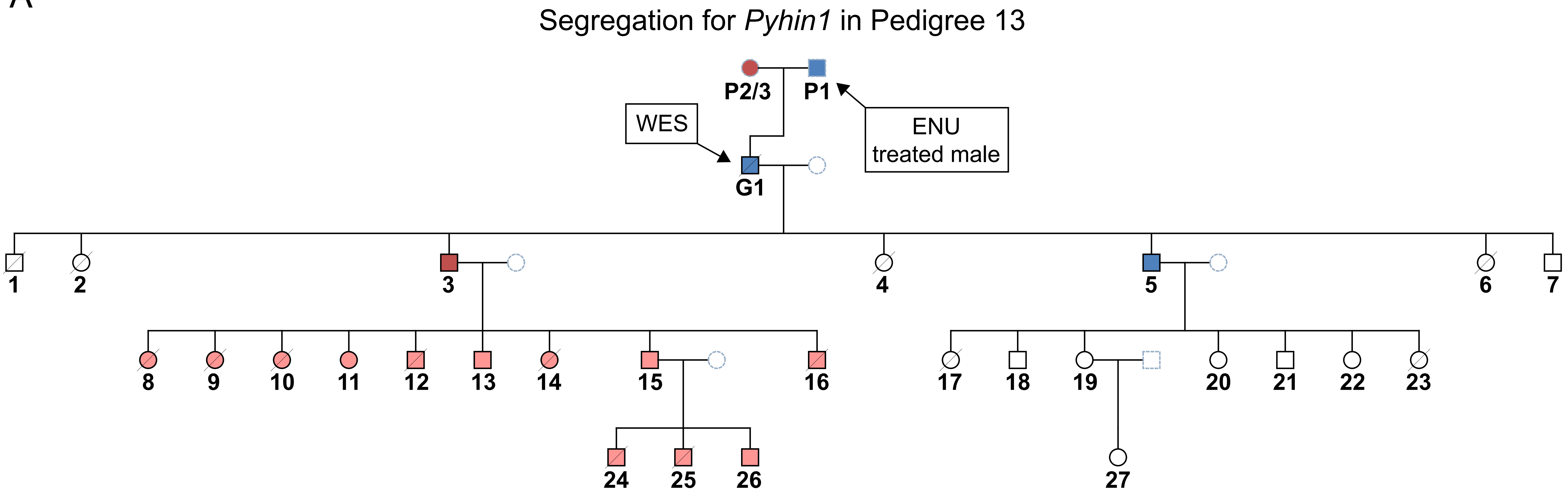

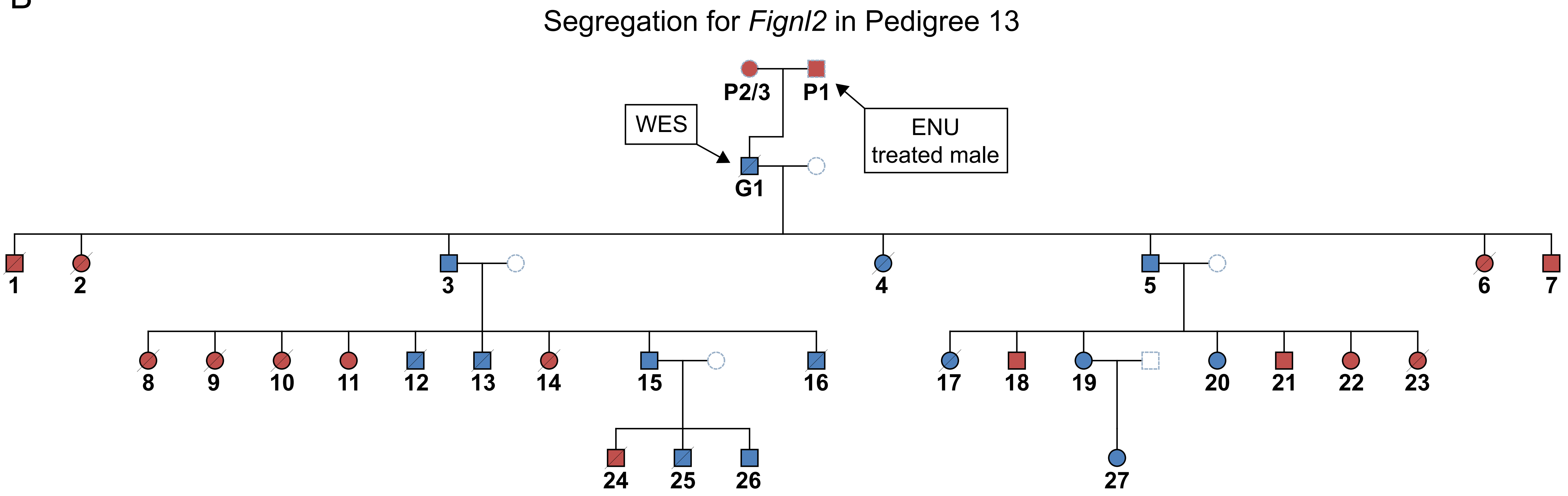

### Supplementary Materials

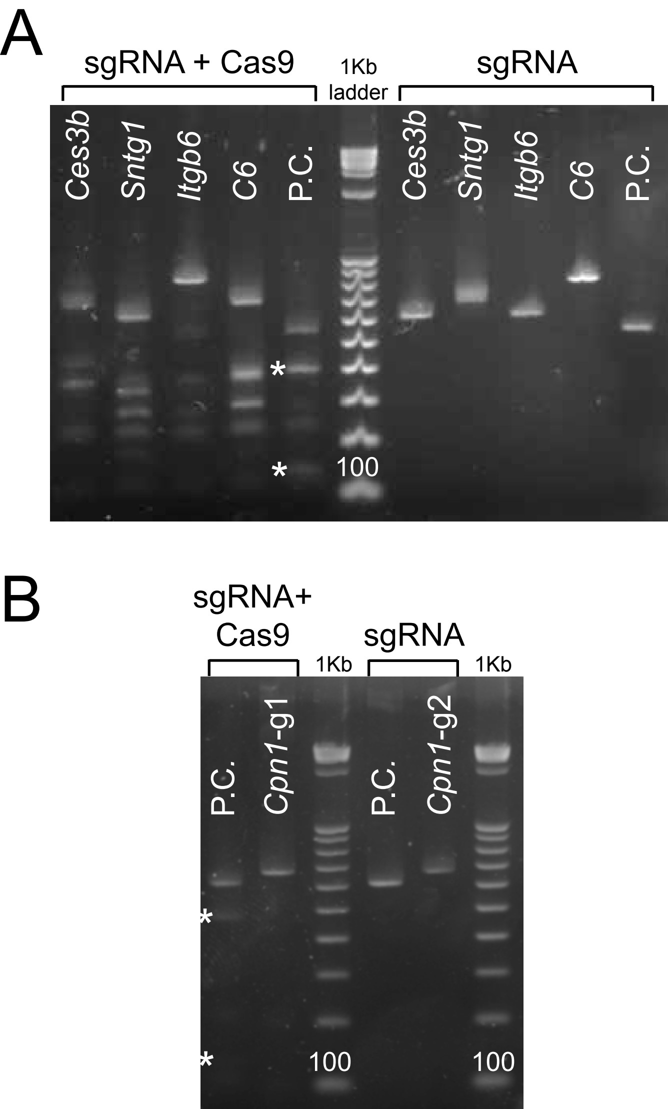
